# AMPK modulates a DEAH Box RNA-helicase to attenuate TOR signaling and establish developmental quiescence

**DOI:** 10.1101/2025.04.03.646977

**Authors:** Sabih Rashid, Richard Roy

## Abstract

Developmental plasticity allows organisms to adapt to environmental stress and improve reproductive fitness. *Caenorhabditis elegans* adapts to starvation and other stressors by transiting through an alternate developmental stage called dauer, which allows them to remain quiescent for several months, and yet fully retain reproductive fitness when they resume development. The AMP-activated protein kinase (AMPK) is essential for this plasticity as its compromise leads to germline hyperplasia during the dauer stage, dramatically reducing post-dauer fertility upon recovery from this stage, while also shortening survival.

We identified a putative RNA-binding helicase (HZL-1) that is targeted by AMPK, the compromise of which suppresses several AMPK mutant phenotypes. HZL-1 shares significant similarity with the conserved HELZ family of RNA helicases, possessing characteristic DEAH helicase motifs, a predicted ATP binding motif, and three intrinsically disordered regions that are crucial for its localization and function. Curiously, HZL-1 is expressed and exerts its function in the intestine, yet its elimination suppresses the aberrant germ cell proliferation, while restoring germline quiescence and subsequent post-dauer fertility. CLIP-seq data revealed that HZL-1 binds several mRNAs during the dauer stage, resulting in a pronounced germline hyperplasia in the dauer germ line of AMPK mutants. Among these, the most enriched RNA bound to HZL-1, *argk-1*, is required for fertility in HZL-1 mutants, and functions by suppressing TOR activity in the germ line of AMPK dauer larvae, thereby preserving germline quiescence. These findings underscore the intricate role of RNAs and RNA-binding helicases in the complex interplay of genetic signals that animals have acquired to ensure their effective transit through periods of environmental challenge.

## Introduction

Environmental fluctuations pose important challenges to all organisms, often limiting access to resources required for growth and reproduction. Through adaptation to these changes, some species have evolved a means of altering their developmental trajectory in response to environmental or physiological stressors [1]. This “developmental plasticity” often involves periods of punctuated cell cycle quiescence [2], and in some cases, the execution of alternate developmental stages [3–4], The decisions to execute one choice versus another are usually very tightly regulated by genetic pathways that respond to specific environmental and/or developmental cues to ensure the optimal developmental trajectory that will maximize fitness is robustly selected. One of the best-studied models for developmental plasticity is the nematode *Caenorhabditis elegans*, which exhibits numerous forms of plasticity, from the L1 larval stage diapause to the distinct dauer diapause, and even an ability to undergo arrest as reproductive adults [5–9]

The dauer diapause is of particular interest since many of the genetic mutations that alter this developmental decision are also involved in tissue growth, scaling, and longevity [10–12]. *C. elegans* larvae adopt this alternative developmental stage in response to suboptimal environmental conditions, including lack of nutrients, elevated temperature, and high population density. Transition into this stage permits the animals to survive these challenges until growth conditions improve [13]. These larvae are characterized by global developmental quiescence, altered metabolism, a modified protective cuticle, and widespread changes in gene expression [14]. Moreover, these developmental changes allow the animals to survive for up to four months without feeding, while the average lifespan of a non-dauer *C. elegans* is approximately two to three weeks.

Remarkably, dauer larvae can quickly readjust their metabolism and resume their growth when environmental conditions improve, allowing them to return to a reproductive developmental trajectory. Furthermore, there are no obvious negative repercussions to fitness by passaging through the dauer stage, as animals remain fertile and have similar longevity to counterparts that do not enter the diapause state. However, post-dauer *C. elegans* do have an altered life history that is recorded in their chromatin, like a molecular memory, as well as slight changes in their reproductive capabilities, both of which persist transgenerationally [15–16].

A large number of signaling pathways must work in concert, not only to induce the changes required for the transition into and out of dauer, but also to preserve the integrity of *C. elegans* cellular functions and ensure animals are fit and fertile following recovery from the dauer stage. Cellular pathways that otherwise promote cell proliferation and growth must respond to changes associated with dauer onset [17]. The signaling pathways that drive reproductive growth are attenuated, while the change in energy levels that is associated with the diapause state results in the activation of AMP-activated protein kinase (AMPK). Conversely, the target of Rapamycin (TOR) signaling pathway has also been implicated in the formation of dauer larvae, as loss of TOR components results in the induction of dauer-like arrest, with corresponding changes in metabolism [18]. During the dauer stage, when nutrients are less abundant, TOR acts as a key signaling molecule that informs the *C. elegans* larva whether to adopt a reproductive mode of development, or to execute the diapause and to forego development until environmental conditions improve.

AMPK plays a significant role in establishing and maintaining germline quiescence during dauer. Previously, it was observed that loss of AMPK leads to germline hyperplasia during the dauer stage. Importantly, in addition to the appearance of various somatic defects in the post-dauer animals, most of the dauer larvae that do recover become sterile [19]. This is associated with a general misregulation of germline chromatin modifications, which has a significant impact on the gene expression of these animals, and may indeed contribute to the phenotypic changes characteristic of AMPK mutant dauer and post-dauer larvae.

Curiously, disabling the activity of some of the critical genes involved in the RNA interference pathway such as *dcr-1,* or the Argonaute protein *ergo-1,* partially suppressed the germline hyperplasia and post-dauer sterility [19]. Therefore, AMPK must be involved in regulating appropriate small RNA homeostasis, as animals lacking AMPK have misregulated small RNAs in the dauer stage, suggesting they play a key role in regulating germline quiescence. However, the precise mechanisms by which AMPK influences RNA levels or function in order to preserve germline integrity are presently unclear.

Here, we show that a previously uncharacterized RNA-binding protein acts as a downstream target of AMPK that binds and inhibits RNAs that are critical for germline integrity during the dauer stage. Structural analysis suggests that it is related to DEAD/DEAH-box RNA helicases found in other species, such as HELZ [20–21], and as such we have named it <u>H</u>EL<u>Z</u>-like 1, or HZL-1. Like many other members of this protein family, HZL-1 possesses three intrinsically disordered regions that contribute to its function. As a putative phosphorylation target of AMPK, this protein functions in the intestine to regulate several target mRNAs, most notably the arginine kinase *argk-1*, ultimately impinging on the TOR signaling pathway. In the absence of AMPK signaling, HZL-1 binds to and inhibits the expression of its target RNAs, thereby preventing the required quiescence in the germ line of dauer larvae and promoting a proliferative signal through the activation of TOR. Our findings highlight a role for RNA-binding proteins in enabling the adjustment to nutrient stress by regulating the stability of specific mRNAs.

## Results

### A candidate RNAi survey to identify regulators of post-dauer fertility downstream of AMPK

Previous analyses of *daf-2(e1370)* AMPK-deficient (hereafter simply called *aak(0)*) dauer larvae revealed defects in germline quiescence and post-dauer fertility, some of which were suppressed by compromising specific RNA interference pathway components [19]. This suggests that the loss of AMPK resulted in the misregulation of small RNAs, although it was unclear how AMPK could impact small RNA levels, and how their putative target mRNAs could affect quiescence. To identify potential downstream targets of AMPK that regulate RNAs, we performed a bioinformatic analysis of the *C. elegans* protein sequences that harbored canonical AMPK consensus sites [22], which might indicate they are direct targets of AMPK. We further narrowed the list of targets by focusing on proteins predicted to be associated with small RNAs [23–24].

To determine if any of these putative targets played a functional role in regulating dauer germline quiescence, we performed a post-dauer fertility assay following RNAi against the target genes, reasoning that a suppression of the post-dauer sterility seen in AMPK mutants could indicate the target was abnormally active in AMPK-deficient animals and contributed to the reproductive defect. This screen identified two targets that, when knocked down, resulted in a suppression of post-dauer fertility in AMPK animals: *parp-2*, a poly-ADP ribose polymerase, and an uncharacterized gene, C44H9.4 (S1A Fig).

**S1 Fig. RNAi of putative AMPK targets *parp-2* and *hzl-1* suppresses post-dauer sterility in *aak (0)* animals.**

(A) Post-dauer fertility in *daf-2* and *daf-2;aak(0)* animals following RNAi against putative AMPK phosphorylation target genes, as predicted by GPS 6.0 software.

(B) Post-dauer fertility of control, *aak(0)* and *C44H9.4(0); aak(0)* animals treated with *parp-2* RNAi. L4440 serves as the empty vector control.

Post-dauer fertility data represents three independent trials, with the mean represented by columns and values for individual trials indicated by small circles. n=50 for each trial. ****p < 0.0001 using two-way ANOVA for the indicated comparisons.

A genetic deletion mutant of C44H9.4, *ok1688,* displayed a similar suppression of post-dauer sterility (Fig 1A). Conversely, RNAi against *parp-2* in this background did not result in additive effects on fertility (S1B Fig), suggesting the two genes may be acting in a common linear pathway. Loss of either gene in control dauer backgrounds (*daf-2)* had no effect on fertility, nor did it cause any observable phenotypes. We chose to focus on C44H9.4 for further study due to its novel nature, as well as its putative roles in RNA regulation.

**Fig 1.**
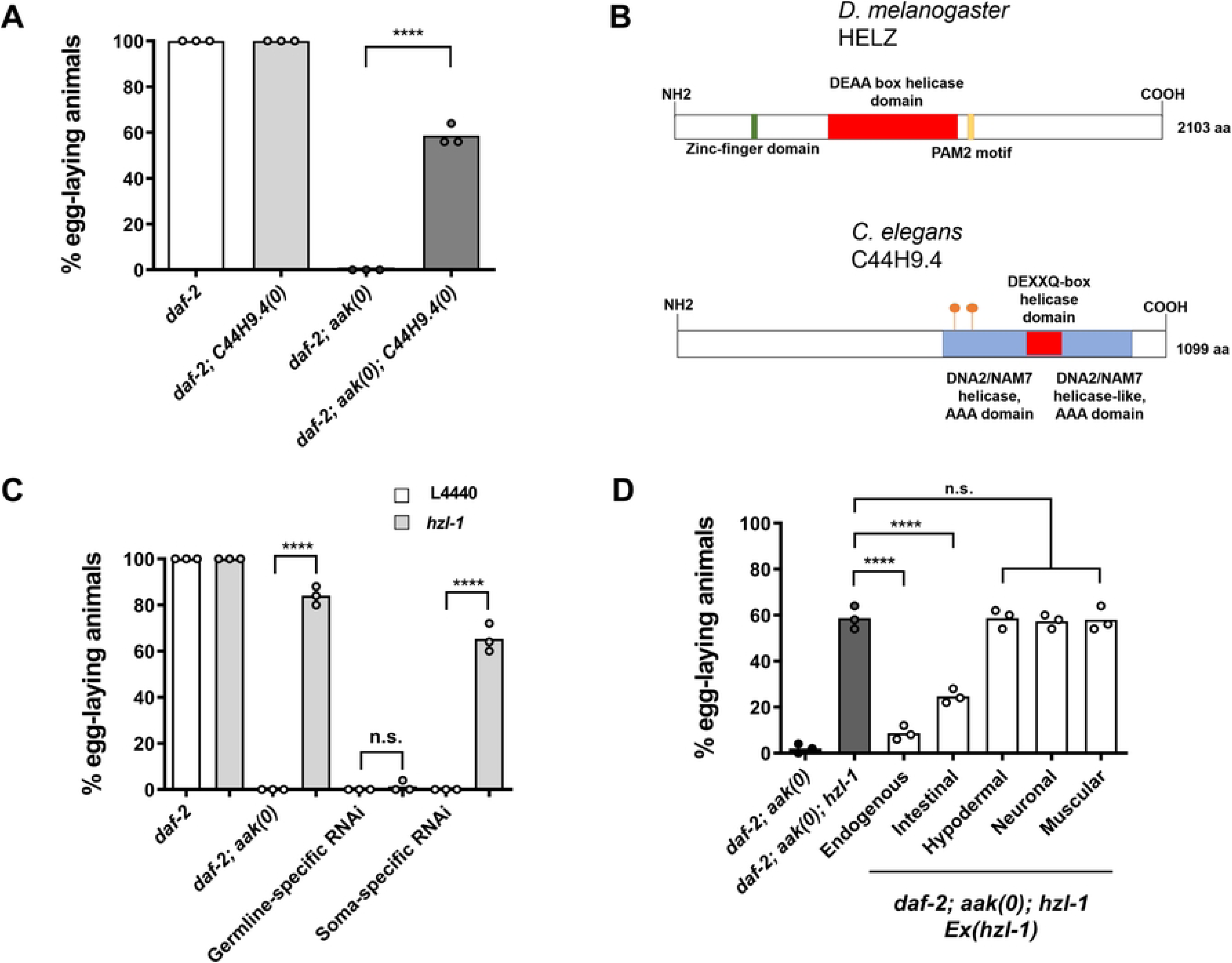
Loss of HZL-1, a putative RNA-binding helicase, suppresses post-dauer sterility in *aak(0)* mutants. (A) Quantification of post-dauer fertility of *daf-2*, *aak(0)* and C44H9.4/*hzl-1* mutants. (B) Schematic diagram depicting (top) the protein HELZ, a *Drosophila melanogaster* ortholog of C44H9.4 based on BLASTP analysis and (bottom) *C. elegans* C44H9.4. Depicted RNA helicase domains predicted by InterPro. Putative AMPK phosphorylation sites at S588 and S636, as predicted by GPS 6.0 software, are shown as orange beacons. (C) Quantification of post-dauer fertility in animals following tissue-specific RNAi of C44H9.4/*hzl-1* in *daf-2; aak(0)* animals. Whole animal RNAi (*daf-2; aak(0)*), soma-specific RNAi (*daf-2; aak(0); rde-1; sur-5p::rde-1*), and germline-specific RNAi (*daf-2; aak(0); rde-1; sun-1p::rde-1*) was performed. (D) Quantification of post-dauer fertility in *hzl-1* mutant animals rescued by a wild-type copy of *hzl-1*. Transgenic insertion of a wild-type copy of *hzl-1* was done in these mutants under the control of its endogenous promoter, or under an intestinal (*nhx-2*), hypodermal (*wrt-2*), neuronal (*rgef-1*), or muscular (*myo-3*) tissue-specific promoters. All post-dauer fertility data represent three independent trials, with the mean represented by columns and individual values by small circles. n=50 for each trial. ****p < 0.0001 using one-way ANOVA for the indicated comparisons, two-way ANOVA for (C).

Structural predictions of the C44H9.4 protein indicate that it contains DNA2/NAM7 helicase domains, as well as a DEXXQ-box helicase domain (Fig 1B). Furthermore, sequence comparisons indicated that the predicted C44H9.4 proteins is an orthologue of an RNA-binding helicase called HELZ, that has been characterized in *Homo sapiens* and *Drosophila melanogaster* [20–21]. Notably, C44H9.4 lacks the Zinc-finger domain and PAM2 motif present in HELZ, suggesting that its function may have diverged, despite having retained the RNA-binding motifs. Nevertheless, due to the shared similarities between these proteins, we will hereafter refer to it as <u>H</u>EL<u>Z</u>-like HZL-1 or *hzl-1*.

Given that the loss of *hzl-1* was sufficient to restore fertility in post-dauer AMPK mutants, we questioned whether its function was required autonomously in the germ line *per se*, or if it acted non-cell autonomously similar to AMPK, being required in neurons or other somatic tissues [19]. Tissue-specific RNAi performed against *hzl-1* in the germ line revealed no effects on fertility, whereas soma-specific RNAi was sufficient to suppress sterility nearly to the same levels as whole-animal RNAi (Fig 1C), hinting that the protein may function in tissues other than the germ cells but communicates information to the germ line. To identify which tissues *hzl-1* may be active in, we inserted a transgenic wild-type copy of the protein into the mutant strain under its endogenous promoter, as well as various tissue-specific promoters. Expression of *hzl-1* under its own endogenous promoter led to near-complete reversion of the suppression of post-dauer sterility, as animals were observed to be mostly sterile, suggesting the gene’s function had been restored (Fig 1D). In addition, intestinally-expressed *hzl-1* was able to partially revert the suppression as well, with significantly more of these transgenic animals being post-dauer sterile compared to the mutant strain, whereas expression under other tissue-specific promoters had no effect. These data therefore suggest that HZL-1 functions in the intestine of AMPK mutant larvae to mediate its effects on germline quiescence during the dauer stage.

### A putative AMPK phosphorylation site within HZL-1 is required for its function

HZL-1 function is abnormally active in AMPK-deficient dauer animals, and so we hypothesized that it may be targeted for inhibition by AMPK, likely through phosphorylation. HZL-1 has two predicted AMPK phosphorylation motifs in its amino acid sequence. Phosphorylation by AMPK can result in the inhibition of proteins through various means, including the induction of 14-3-3 binding to the target [25]. To determine whether AMPK regulates HZL-1 activity by phosphorylation, we mutated the predicted phosphoacceptor sites on the protein.

Phosphomimetic variants were created by replacing the acceptor serine with an aspartic acid (Fig 2A). Conversely, non-phosphorylable variants were created by replacing the same serine residue with a neutral alanine. Variants were also generated wherein both predicted sites were mutated to either phosphomimetic or non-phosphorylable residues. We reasoned that a phosphomimetic variant of HZL-1 would mimic phosphorylation by AMPK, meaning that its activity would presumably be attenuated and thus function similarly to the deletion mutant of HZL-1, resulting in fertile post-dauer adults. Conversely, a non-phosphorylable variant might result in HZL-1 being aberrantly active in the wild-type, since such a protein would no longer be regulated by AMPK, and thus might escape from with germline quiescence in the dauer stage, despite the presence of active AMPK.

**Fig 2.**
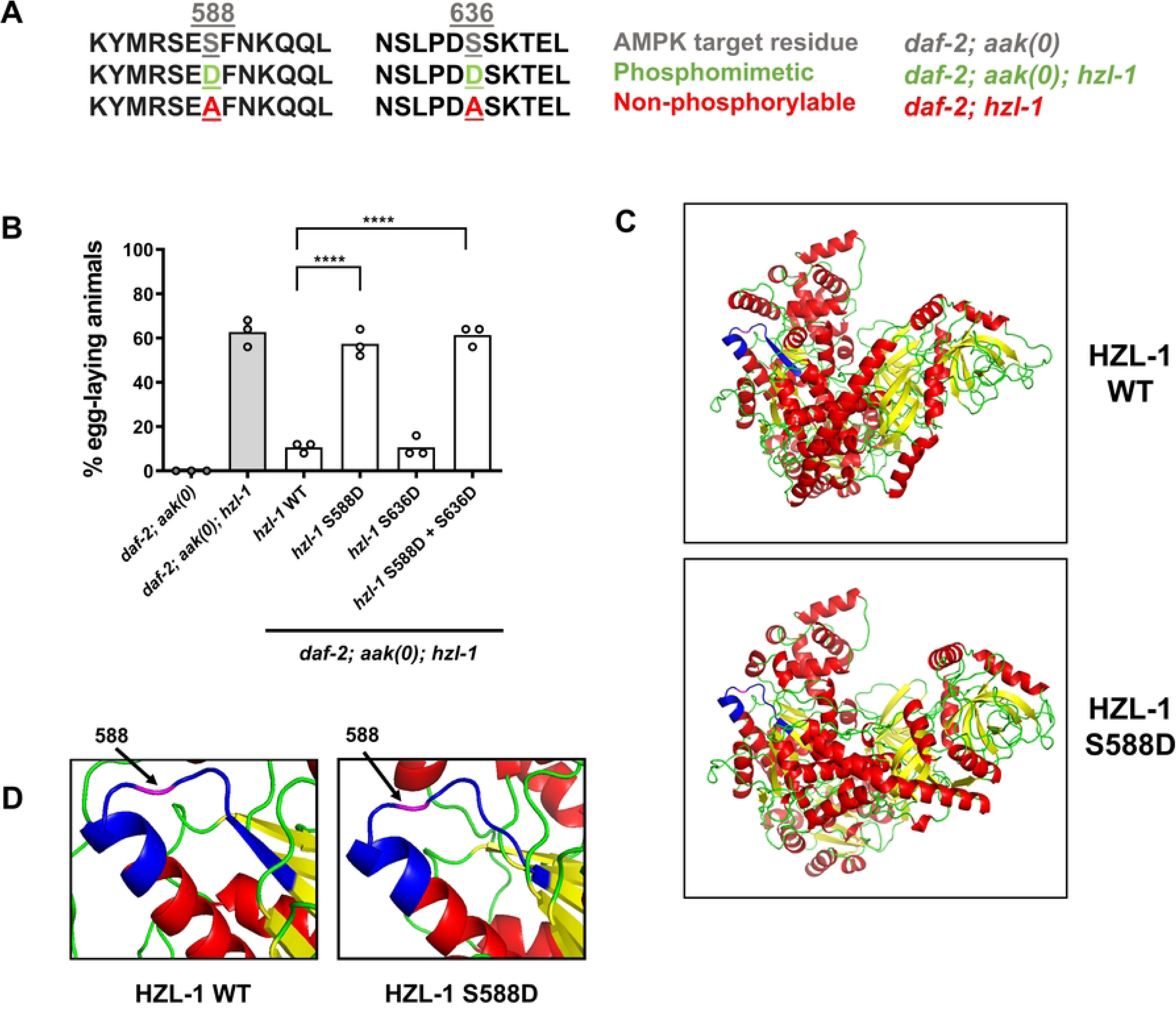
A putative AMPK phosphorylation site on HZL-1 modulates its function. (A) Amino acid sequence of the predicted AMPK phosphorylation site at serine 588 or serine 636 in HZL-1 including indicated phosphomimetic/phosphonull (non-phosphorylable) changes (**D** and **A** in green and red, respectively). The genetic background that each phosphovariant gene was inserted into is listed on the right. (B) Post-dauer fertility of control animals, or those harboring transgenes that express phosphomimetic HZL-1 variants. Wild-type or mutated *hzl-1* was transgenically inserted into *daf-2; aak(0); hzl-1* strains. (C) Predicted protein models of HZL-1 wild-type or S588D phosphomimetic mutant, as generated by Alphafold 3 [29]. Purple indicates the residue at position 588 and blue denotes the region around it. (D) Closeup of HZL-1 wild-type and S588D mutant proteins from C), focused around the 588 residue and altered loop structure. Purple indicates the residue at position 588 and blue denotes the region around it.

When assessing the post-dauer fertility of these variants, we noted that the phosphomimetic strain with the S588 site mutated was indeed post-dauer fertile despite it lacking all AMPK signaling, very similar to animals that lacked *hzl-1* (Fig 2B). This suggests that AMPK could potentially phosphorylate HZL-1 at S588, resulting in its inhibition. However, the independent mutation of the second site at S636 appeared to have no effect, as transgenic animals harboring this phospho-site variant exhibited similar levels of fertility to the intact HZL-1 transgene control. There appeared to be no additive effect of mutating both sites, suggesting that S588 is the only phosphorylable residue that is functionally relevant.

When assessing the fertility of the non-phosphorylable variants in a *daf-2; hzl-1* background, we observed no changes in fertility, as all animals were fully fertile similar to the control. Furthermore, mutation of both sites had no effect on any of the observed phenotypes (S2A Fig). This indicates that even if HZL-1 is active during the dauer stage due to a lack of phosphorylation by AMPK, its active state may not be sufficient alone to interfere with germline integrity. Alternatively, other targets that act downstream of AMPK signaling might counteract the function of HZL-1, either by inhibiting the protein itself, or by promoting a parallel pro-quiescent signal that circumvents any effect of HZL-1 [25]

**S2 Fig. Phosphorylation at S588 attenuates HZL-1 function through mechanisms independent of differential degradation or localization**

(A) Post-dauer fertility of animals in control genetic backgrounds, or transgenics variants with non-phosphorylable mutations in the predicted AMPK consensus sites present in HZL-1. Wild-type or mutated *hzl-1* was introduced into *daf-2; hzl-1* strains as extrachromosomal arrays.

All post-dauer fertility data represent three independent trials, with the mean represented by columns and values for individual trials indicated by small circles. n=50 for each trial. ****p < 0.0001, ***p < 0.001 using one-way ANOVA for the indicated comparisons.

(B) Levels of HZL-1::GFP detected by Western blot using anti-GFP antibodies. Western analysis done with phosphomimetic HZL-1 mutants compared with wild type (WT). α-tubulin (Bottom) is the loading control.

Phosphorylation by AMPK can inhibit proteins in several different ways, including degradation of the protein, induction of conformation changes, or sequestering to different cellular compartments [26–27]. Many AMPK targets specifically are sequestered and degraded through generation of 14-3-3 target sites following phosphorylation [26, 28]. To identify whether phosphorylation by AMPK results in degradation of HZL-1, we performed Western blot against the GFP tagged to HZL-1 in our phosphor-variants. Protein levels remained largely identical in all of our phosphomimetic variants, including the S588D variant, suggesting there is no phospho-dependent degradation of HZL-1 (S2B-C Fig).

We then wanted to know whether the localization of the protein is affected by phosphorylation. It is difficult to image *hzl-1* due to the low endogenous levels of expression, and therefore we used the intestinally-expressed variant, which we showed can restore fertility in the post-dauer mutants (Fig 1D). Expression of the GFP tag is strong in the intestine (S2D Fig). Furthermore, when we express the S588 phosphomimetic under the intestinal promoter, we see the same expression, suggesting that the additional charge conferred by the aspartate on the phosphomimetic variant does not result in any change in the cellular localization of HZL-1, and the protein’s function may instead be inhibited via some other mechanism, such as a conformational change.

We looked at the predicted structures of both the wild-type and the S588D phosphomimetic variant of HZL-1 using AlphaFold [29]. There is no obvious difference in the larger structure of the two proteins when compared side by side (Fig 2C). However, when focusing on the 588 residue, we note that there appears to be a larger space in the S588D variant compared to wild-type around that residue (Fig 2D). If HZL-1 is indeed an RNA-binding helicase, this difference in structure may impact the ability of the protein to bind RNA, as both the electrostatic charge as well as physical structure impacts interactions between RNA and protein [30–31]. Phosphorylation by AMPK may thus induce this physical change in the structure of HZL-1, impacting its ability to bind RNAs without affecting its expression or localization.

### Intrinsically disordered regions in HZL-1 are required for its expression during dauer and its regulation of post-dauer fertility

Many RNA-binding helicases are able to form liquid-liquid condensates that facilitate further interactions with RNAs or other proteins [32]. Intrinsically disordered regions (IDRs) within the protein can confer the ability to form these condensates and are frequently observed in RNA helicases [33–34]. Given the predicted domains on HZL-1 indicate that it could be an RNA-binding helicase, we wondered whether the protein also harbored IDRs, and whether they might contribute to its function, potentially through the formation of condensates. Analysis of the amino acid sequence using the resource IUPred3 [35] revealed that HZL-1 possessed three predicted IDRs (Fig 3A-B).

**Fig 3.**
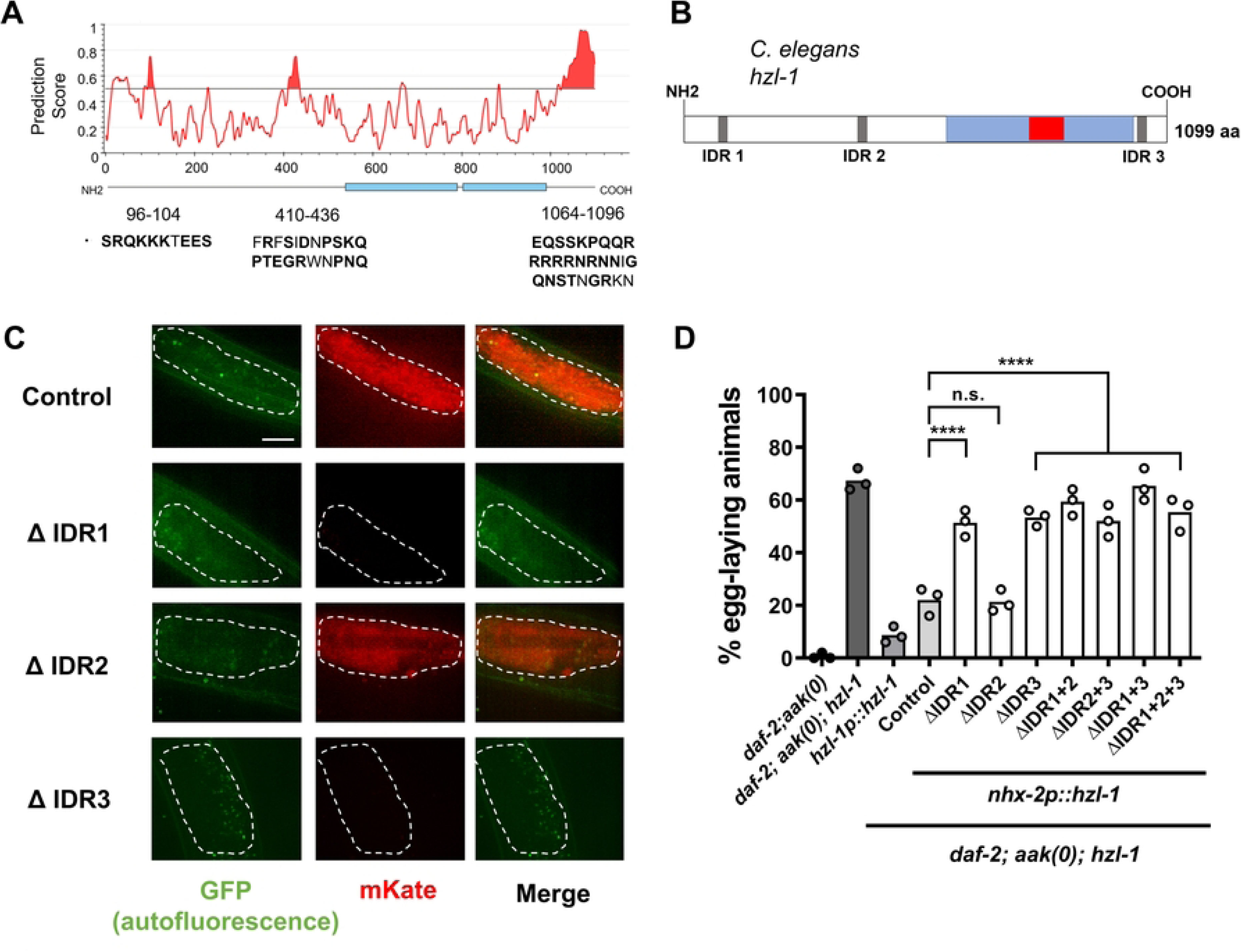
HZL-1 possesses three predicted IDRs that are differentially required for its expression and function. (A) Intrinsically disordered domains present in the HZL-1 amino acid sequence as predicted by IUPred3. Residues that correspond to the predicted peaks are indicated below the graph in capital letters, with disorder-contributing residues in bold. (B) Updated protein schematic of HZL-1 with location of predicted IDRs indicated. (C) Confocal images of HZL-1 expressed under the intestinal promoter (*nhx-2*) in *aak(0); hzl-1* mutants. Dauer larvae (Control) were all *daf-2* maintained at 25°C. Each micrograph shows the posterior intestine that expresses transgenic HZL-1 variants with deletions of individual IDR sequences (IDR1, IDR2 and IDR3). Scale bar = 10 µm (D) Post-dauer fertility of *daf-2; aak(0); hzl-1* rescued with *nhx-2p*:: *hzl-1* with deletion variants of single or compound IDR deletions (ΔIDR1, ΔIDR3 and ΔIDR3) from the HZL-1 sequence. ‘Control’ refers to intact wild-type HZL-1 expressed in the gut under the control of the *nhx-2* promoter. Post-dauer fertility data represents the results from three independent trials, where the mean is represented by columns and values for individual trials indicated by small circles. n=50 for each trial.****p < 0.0001 using one-way ANOVA for the indicated comparisons.

To further investigate whether these IDRs contribute to the function of HZL-1, we designed a series of transgenic variants with single or compound deletions in one or more of the predicted IDRs and expressed them in the *hzl-1* null mutant (Fig 1D). When HZL-1 is expressed under the control of its endogenous promoter it is difficult to detect, potentially due to low expression. Therefore, in order to determine the role of these IDRs in the function of HZL-1, we expressed the IDR variants using a strong intestinal promoter that is sufficient to partially rescue the null mutant when driving the wild-type version of HZL-1 (Fig 1D). Using an mKate reporter, we could detect strong fluorescent signal in the intestine of dauer larvae when we drove the expression of an intact (i.e. wild-type) HZL-1 protein (Fig 3C, top row). In contrast to the wild-type protein, the IDR1 or IDR3 deletions were undetectable during the dauer stage (Fig 3C, rows 2 and 4), whereas the IDR2 deletion had no effect, as fluorescence was similar to the wild-type control. To confirm that deletion of the IDRs did not affect the expression of *hzl-1*, we performed qPCR to measure levels of *hzl-1* in our IDR deletion strains, and found that the levels of mRNA were consistent regardless of whether wild-type or IDR variants of *hzl-1* were expressed (S3 Fig).

**S3 Fig. Removal of IDR sequences does not affect *hzl-1* mRNA levels**

qPCR quantification of *hzl-1* mRNA levels in dauer *daf-2; aak(0); hzl-1* mutants that harbored variants of the reverting transgene *nhx-2p::hzl-1,* that had one or more IDR sequences deleted from the *hzl-1* sequence. Statistical significance was tested using one-way ANOVA.

Furthermore, analysis of the post-dauer fertility of the ΔIDR transgenic strains revealed that loss of IDR1 or 3 increased their fertility compared to the control transgenic strain expressing wild-type HZL-1 (Fig 3D). This suggests that these variants of the protein impede the function of HZL-1 such that unlike its wild-type counterpart, even though the protein is expressed, it does not disrupt germline quiescence because it is not functional. These findings are consistent with the observed changes in protein expression, further corroborating the requirement of IDR1 and IDR3 for the proper activity, or stability of HZL-1 during the dauer stage. Furthermore, the significance of IDRs within HZL-1 provides further evidence that it may be an RNA-binding helicase, as IDRs are often necessary for helicases that regulate RNAs [36].

### Transcriptomic analysis of AMPK and HZL-1 mutants reveals widespread gene expression differences primarily in the germ line

The structure and the conserved domains of HZL-1 are consistent with it acting as an RNA-binding protein that may inappropriately affect the levels of growth- or proliferation-specific RNAs in dauer larvae when AMPK activity is compromised. To better understand what changes in RNA homeostasis may be induced by the protein, we performed a transcriptomic analysis of the HZL-1 deletion mutant. We obtained total RNA from dauer and post-dauer animals that were of four different genetic backgrounds: *daf-2* (wild-type control); *daf-2; aak(0)* (AMPK mutant), *daf-2; hzl-1* (HZL-1 mutant control), and *daf-2; aak(0); hzl-1* (HZL-1 and AMPK double mutant). These samples were then submitted for mRNA sequencing, and bioinformatic analysis was subsequently performed (S4A Fig).

**S4 Fig. Protocol for our RNA-Seq.**

(A) Protocol of RNA-Seq methodology. Animals of the indicated genotypes were grown in large quantities at 25°C until reaching day 2 of the dauer stage, and were either harvested for the dauer sample, or allowed to grow at 15 °C for another 4 days before harvesting for post-dauer samples. Total RNA was obtained through Trizol extraction before being sent to a sequencing facility. mRNA sequencing was performed, followed by reference genome assembly. DESeq2 was used for bioinformatic analysis.

(B) Volcano plots depicting the spread of gene expression changes between *aak(0); hzl-1 and aak(0)* dauer and post-dauer (PD) animals

Transcriptional analysis of these mutants revealed widespread gene expression changes during both the dauer and post-dauer stages (S4B Fig). We noted a large number of differentially expressed genes when comparing *aak(0)* to control *daf-2* animals, and comparing *aak(0); hzl-1* animals to *aak(0)* samples. We then performed GO enrichment of our transcriptomic comparisons, and identified reproductive and germ line genes as being the most enriched in the gene set that had increased expression *aak(0)* dauer animals compared to controls (Fig 4A). Notably, this pattern of expression was inverted in the post-dauer comparison, suggesting reproductive genes are expressed at relatively lower levels in AMPK mutant post-dauer adult animals, potentially because their reproductive developmental program was prematurely activated during the dauer stage.

**Fig 4.**
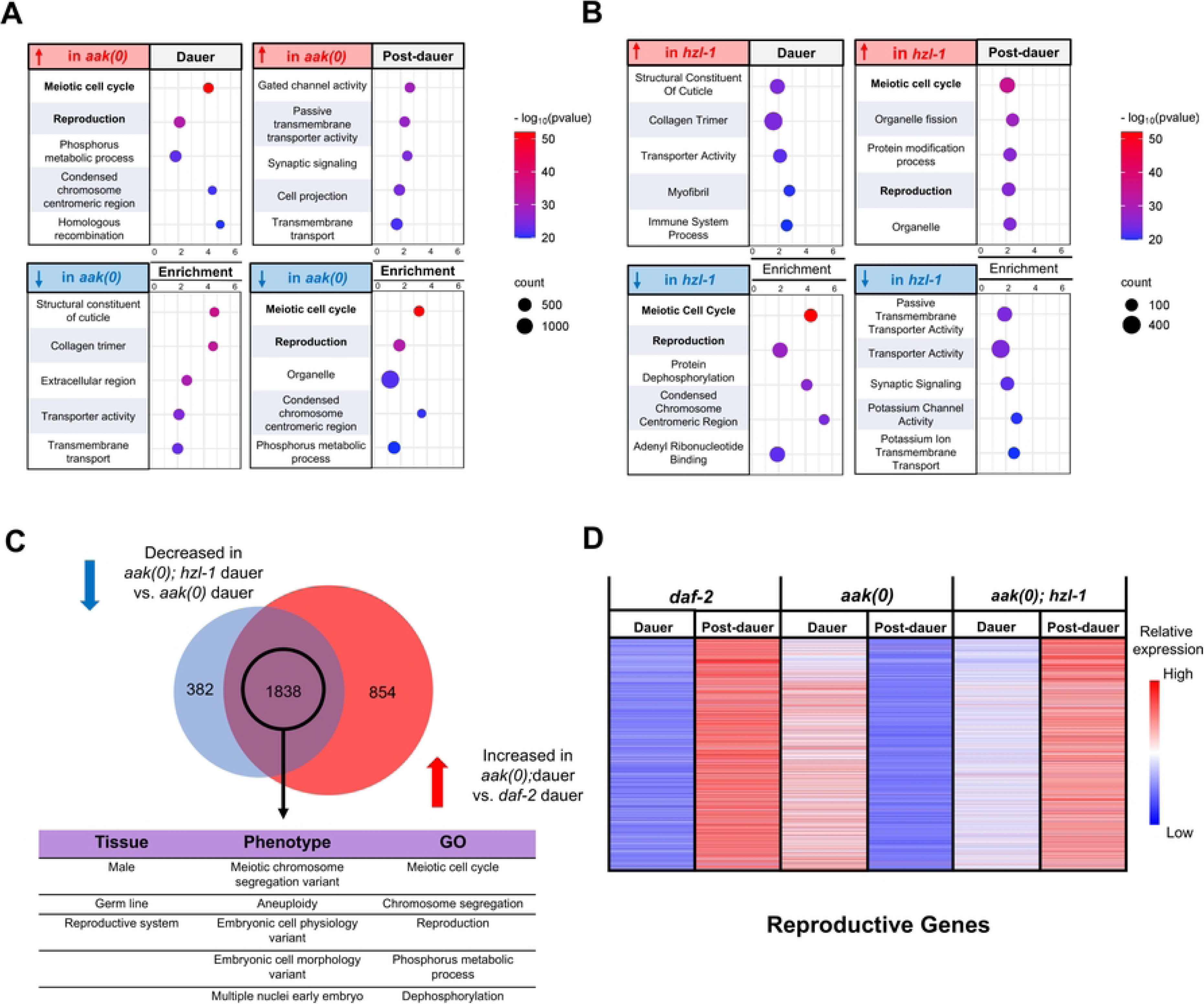
RNA-Seq of AMPK, HZL-1 mutants reveals widespread gene expression differences largely in the germ line. (A) and (B) Bubble plots depicting most enriched GO terms in *aak(0)* animals compared to *daf-2* controls (A), and *hzl-1(0); aak(0)* compared to *aak(0)* (B), ranked by significance based on p-value. A) Left and right graphs represent dauer and post-dauer comparisons, respectively. Top graphs depict genes increased in *daf-2; aak(0)* mutants compared to *daf-2* controls, while bottom graphs depict those genes that were decreased. Size of bubbles indicate number of genes in their respective categories that were enriched in the dataset. GO enrichment was conducted using the Wormbase Gene Set Enrichment Analysis [37–38]. (C) (Top) Venn diagram comparing genes with decreased expression in *hzl-1; aak(0)* dauer vs. *aak(0)* dauer, and genes with increased expression in *daf-2; aak(0)* dauer vs. *daf-2* dauer, indicating a large overlap of 1838 genes. Bottom: Tissue, phenotype, and gene ontology enrichment of overlapped genes, highlighting an enrichment in genes associated with the germ line and reproductive phenotypes. (D) Heatmap of the expression of ∼2000 reproductive genes from (C). This subset of genes is differentially expressed in *aak(0)* and *hzl-1; aak(0)* mutants, as well as in dauer and post-dauer animals. Data represents normalized transcripts per million values for each gene, with higher relative expression in red and lower in blue.

Loss of AMPK thus appears to induce reproductive development aberrantly, even in the dauer stage, where animals are expected to be quiescent. This correlates with our phenotypic analysis of these mutants, which reveals germline hyperplasia and premature spermatogenesis that occurs during the dauer stage [16].

We noted a similar inverse pattern of expression when comparing *aak(0); hzl-1* mutants to *aak(0)* (Fig 4B). Specifically, the GO enrichment in the subset that was decreased in *hzl-1* mutant dauer animals is comparable to the subset that is increased in AMPK mutant dauer data set. A similar pattern is seen when observing the post-dauer datasets, where reproductive and germline genes have increased expression in *hzl-1* mutants compared to *aak(0)*. This suggests the loss of *hzl-1* reverses the transcriptomic changes induced by the absence of AMPK, consistent with our findings that mutation of *hzl-1* partially restores fertility to these animals.

We further compared the subset of genes with increased expression in *aak(0)* dauer vs controls, with those with decreased expression in the suppressed AMPK mutant dauer larvae (*aak(0); hzl-1)* vs the non-suppressed animals (*aak(0))*, and identified a significant overlap (Fig 4C). Enrichment analysis of this shared gene set suggests they are, indeed, primarily reproductive genes. When comparing expression of these reproductive genes across our datasets, it appears that the *aak(0)* dauer transcriptome is similar to that of *daf-2* control post-dauer, whereas the *aak(0); hzl-1* dauer transcriptome is closer to *daf-2* dauer (Fig 4D). This is true for the post-dauer gene expression of *aak(0); hzl-1* mutants as well. We conclude that HZL-1 likely affects an ensemble of transcripts that are involved in the induction of quiescence to ensure germ cell growth and cell proliferation during larval development, namely during the L3 stage. Under dauer inducing conditions, its activity is attenuated by AMPK-mediated phosphorylation. However, in the *aak(0)* mutant background, HZL-1 is abnormally active in the dauer stage and promotes reproductive development through effects on the transcriptome, contributing to the defects seen in these mutants. As a result, in the *aak(0); hzl-1* mutant background, the transcriptome largely comprises mRNAs that are more typical of a replete state, at least as it pertains to reproductive and germline genes.

### HZL-1 binds mRNAs which may contribute to its regulation of dauer germline quiescence in AMPK mutants

We observed widespread changes in mRNA levels resulting from the loss of *hzl-1*, but it was unclear how this protein was able to have such a major effect on the *C. elegans* transcriptome. Since HZL-1 possesses all the structural features for it to act as an RNA-binding protein, we sought to verify if HZL-1 could indeed bind specific RNAs to modulate their levels, such that when it is misregulated these RNAs might perturb the ability of AMPK mutants to maintain dauer quiescence.

We performed cross-linking immunoprecipitation on HZL-1 followed by RNA sequence analysis (CLIP-seq), by using our intestinally-expressed GFP-tagged HZL-1 strain, which we previously demonstrated was sufficient to restore HZL-1 function (protocol in S5A Fig). Using both *daf-2* control and *daf-2; aak(0)* strains harboring this transgene, we used an anti-GFP antibody to immunoprecipitate RNA targets from a total RNA extract obtained from each genotype. The bound RNAs were then collected and were subjected to RNA-seq and bioinformatic analysis. As a negative control, we also carried out the IP and RNA extraction on a strain expressing GFP under the same intestinal promoter, but without *hzl-1*.

**S5 Fig. Protocol of CLIP-Seq methodology and tissue/phenotype enrichment of CLIP-Seq data**

(A) Protocol of CLIP-Seq methodology. Animals of the indicated genotypes were grown in large quantities at 25°C until they reached day 2 of the dauer stage, and then were subjected to formaldehyde cross-linking, followed by immunoprecipitation with an anti-GFP antibody. Trizol RNA extraction was subsequently performed on the immunoprecipitated proteins. mRNA and small RNA sequencing was performed on the total RNA obtained, followed by reference genome assembly.

(B) Bubble plots depicting Top 100 RNAs bound to HZL-1in the *daf-2* control background, ranked by significance based on p-value. Size of bubbles indicate number of genes in their respective categories that were enriched in the indicated dataset. GO enrichment analysis was conducted using the Wormbase Gene Set Enrichment Analysis [37–38].

The CLIP-seq experiment revealed that numerous RNAs were bound by HZL-1 in both the control (*daf-2)* and *aak(0)* mutant backgrounds. Notably, no small RNAs were detected, and only mRNAs were bound to HZL-1. We compared the data from the *daf-2* and *aak(0)* samples to the negative control (i.e. only GFP) in order to identify any bound targets that may simply be noise or background, and eliminated targets from analysis that were strongly detected in the negative control.

We next sought to determine what types of RNAs were bound to *hzl-1*. Gene enrichment analysis of the bound RNAs revealed a diverse set of gene categories that were present in the CLIP (Fig 5A-B). We analyzed the most enriched RNAs in both the *daf-2* (S5B Fig) and *aak(0)* (Fig 5A) sample, as well as those that had a significant fold change increase when comparing the *aak(0)* sample to our *daf-2* controls (Fig 5B). In the *daf-2* background, we noted enrichment of RNAs associated with a wide variety of functions, including lifespan and development (S5B Fig). The most enriched targets in the *aak(0)* background were associated with metabolism and detoxification (Fig 5A). To our surprise, there was no enrichment of germ line or reproduction associated gene transcripts. This was curious as we showed that HZL-1 exerts its effects on the germ line of dauer animals, but it does not appear to directly regulate many mRNAs of germ line associated genes.

**Fig 5.**
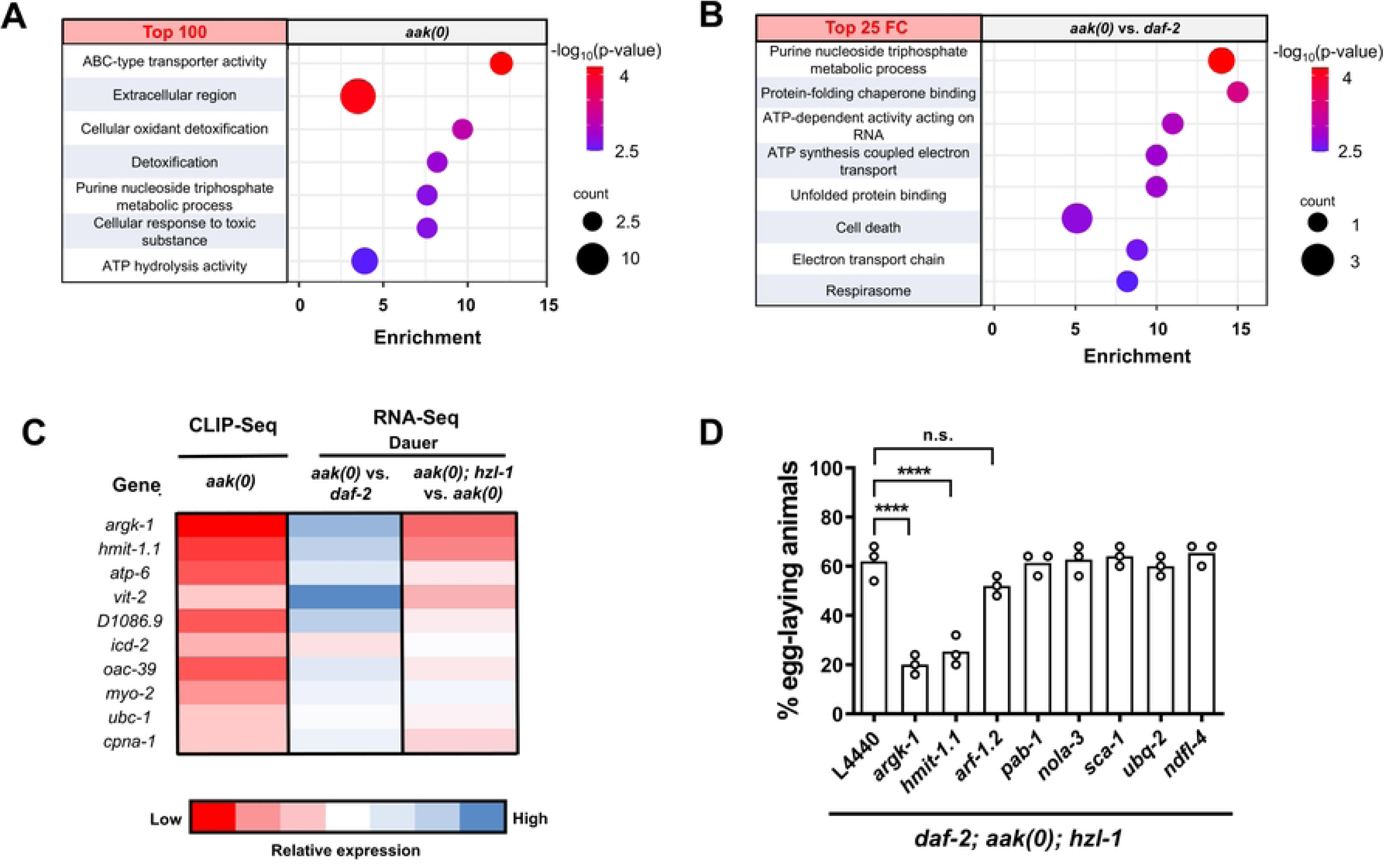
HZL-1 binds numerous RNAs and negatively regulates their expression. (A) and (B) Bubble plots depicting most enriched GO terms in RNAs bound to HZL-1, based on CLIP-Seq data, ranked by significance based on p-value. A) Enrichment of top 100 RNAs bound to HZL-1 in *aak(0)* mutants. B) Top 25 RNAs with biggest fold change increase in the *aak(0)* dataset compared to *daf-2.* Size of bubbles indicate number of genes in their respective categories that were enriched in the indicated dataset. GO enrichment was conducted using the Wormbase Gene Set Enrichment Analysis [37–38]. (C) Heatmap of CLIP-Seq fold change of most highly expressed genes in the *aak(0)* dataset, compared to fold changes of those genes in the *aak(0)* dauer vs. *daf-2* dauer, and *hzl-1(0); aak(0)* dauer vs. *aak(0)* dauer RNA-Seq datasets. Heatmaps generated based on log_2_ fold change of indicated comparisons. (D) Post-dauer fertility of *daf-2; aak(0); hzl-1(0)* animals following RNAi against select targets enriched in CLIP-Seq *aak* dataset. L4440 empty vector serves as a control. Post-dauer fertility data represent the results from three independent trials, where the mean is represented by columns and values for individual trials indicated by small circles. n=50 for each trial. ****p < 0.0001 using one-way ANOVA for the indicated comparisons.

When looking at those RNAs with the biggest fold change difference between our *aak(0)* and *daf-2* samples, we saw an increase in metabolic target enrichment, specifically associated with the electron transport chain and related functions. This may reflect the role of HZL-1 in the intestine, a tissue that responds to diet and nutrition through adjustments to metabolism [39]. Furthermore, in the absence of the metabolic regulator AMPK, perhaps HZL-1 specifically targets RNAs related to energy metabolism. This regulation could potentially influence the metabolic phenotypes that we have previously characterized in AMPK mutant dauer larvae [40].

One notable target of HZL-1 that we obtained in our CLIP-Seq data was *argk-1*, which was the most enriched RNA in our *aak(0)* samples across three biological replicates, and not present at all in the control. ARGK-1 is an arginine kinase that is similar to creatine kinases in mammals, and was previously shown to be enriched in RSKS-1/S6K mutants in *C. elegans*, suggesting it functions downstream of TOR signaling [41]. It is expressed in the intestine, similar to HZL-1, and levels of the protein also positively correlate with AMPK activity, as deletion of *argk-1* was shown to reduce levels of phosphorylated AAK-2. Thus, its prominence in our CLIP-Seq data may indicate that regulation of *argk-1* mRNA by HZL-1 plays a role in the defects we see in *aak(0)* dauer larvae.

To validate that HZL-1 binds *argk-1,* we collected protein extracts from *daf-2* and *aak(0)* transgenic dauer animals, as well as the negative control strain with only GFP. We then performed IP using anti-GFP, with anti-FLAG as a non-specific antibody control (S6A Fig). We performed qPCR against *argk-1* using these samples and observed high levels of the mRNA in the GFP IP samples from the *aak(0)* background relative to input, and this expression was significantly higher than in the non-specific FLAG pulldown sample (S6B Fig). Conversely, no significant *argk-1* levels were detected in the *daf-2* sample or the GFP control.

**S6 Fig. CLIP-qPCR confirms HZL-1 binds *argk-1* RNA in AMPK mutant dauer larvae**

(A) Protocol for CLIP-qPCR. Animals of the indicated genotypes were grown in large quantities at 25 °C until they reached day 2 of the dauer stage, and then were subjected to formaldehyde cross-linking, followed by immunoprecipitation with an anti-GFP antibody. Anti-FLAG was used as a non-specific control. Trizol RNA extraction was subsequently performed on the immunoprecipitated proteins followed by cDNA conversion and qPCR using primers for *argk-1.* S588D refers to the phosphomimetic variant of HZL-1, which was expressed in place of the wild-type variant for that strain. GFP expressed under the intestinal promoter, but without *hzl-1*, served as a negative control.

(B) Quantification of CLIP-qPCR experiment using RNA from either GFP or FLAG IPs. Primers against *argk-1* were used for qPCR. All quantifications were carried out relative to Input i.e. the lysate for the IP, taking into account a dilution factor of 2% i.e. 50 to determine the ‘Adjusted Input’ value. % Input was calculated using the Delta Ct method. *** p < 0.0005 as determined by multiple t-test.

To further confirm that *argk-1* is bound by HZL-1, we also tested the S588 phosphomimetic variant of HZL-1, which we demonstrated has impaired function (Fig 2). Because we showed that the phospho-site does not affect protein abundance, we suspected it might inhibit the RNA-binding function of HZL-1. Consistent with this, our CLIP-qPCR experiment showed that no significant levels of *argk-1* could be detected within the IP pellets isolated from the S588 phosphomimetic variant compared to the FLAG (S6B Fig). We therefore conclude that phosphorylation by AMPK at the S588 residue impairs the RNA-binding capacity of HZL-1, or at least its ability to bind *argk-1* mRNA, perhaps through inducing a conformational change, which has been demonstrated to affect this family of helicases [42].

While the CLIP-Seq analysis suggests that HZL-1 binds many mRNAs, it was unclear how these RNAs were being regulated. Some helicases can act as chaperones, protecting bound RNAs from degradation [43]. Conversely, other helicases can sequester mRNAs, inhibiting their function and/or degrading them directly [44]. We took the list of HZL-1 RNA targets enriched specifically in the *aak(0)* dataset, and compared them to our previous transcriptomic data. We found that many of the bound targets of HZL-1 have decreased abundance in the *aak(0)* background dauer compared to *daf-2* controls (Fig 5C). This pattern is inverted when looking at the *aak(0); hzl-1* dataset i.e. expression of those genes is increased relative to *aak(0).* This suggests that the levels of those RNAs which are bound by wild-type HZL-1 are destabilized in the *aak(0)* mutants. Conversely, those RNAs that are expressed at a higher level in the suppressed *aak(0); hzl-1* mutants compared to *aak(0)* mutants are no longer destabilized by RNA binding by HZL-1, therefore resulting in hyperplasia and post-dauer sterility.

Based on this interpretation, we suspected that the elimination of at least some of these RNAs in the *hzl-1* mutant might revert the suppression, causing the animals to become sterile once more, as the pro-quiescence signal is lost. Indeed, performing individual RNAi against several of the highly enriched CLIP targets, such as *argk-1* and *hmit-1.1,* lead to a reduction in post-dauer fertility of these animals (Fig 5D). We therefore conclude that these two HZL-1 targets, and potentially others, may constitute an ensemble of pro-quiescent signals that affect the dauer germ line, the downstream effects of which are compromised when the transcripts are bound by HZL-1. Notably, we did not observe any effect on fertility by performing RNAi on these targets in a *daf-2* control dauer background. This may indicate that *argk-1* and other targets are not the sole drivers of quiescence, and that other targets regulated by AMPK preserve quiescence in the dauer germ line in the absence of ARGK-1. This finding is consistent with our previous experiments demonstrating that the aberrant activation of HZL-1 through a non-phosphorylable mutant, which would presumably impinge on ARGK-1 activity or abundance, is not sufficient to affect fertility (S2A Fig).

### *argk-1* mediates dauer germline quiescence by influencing TOR activity

We observed that the knockdown of either *argk-1* or *hmit-1.1* resulted in the reversion of suppression of post-dauer sterility in *aak(0); hzl-1* mutants. Why these particular genes affect fertility, however, is unclear. *argk-1* appears to act downstream of S6K and thus TOR signaling [41]. *hmit-1.1* is an H+/myo-inositol transporter, implicated with the osmoprotective response in *C. elegans* [45]. Of the two, we were particularly intrigued by *argk-1* due its role in regulation of metabolism, an aspect that is dramatically altered in AMPK mutants, and is often influenced by the intestine, the tissue where HZL-1 ostensibly functions.

We compared the phenotype of an *argk-1* loss of function mutation with the *argk-1* RNAi, and found that the deletion strain also affected post-dauer fertility in the *aak(0); hzl-1* mutants (Fig 6A), further confirming the influence of ARGK-1 on reproductive capability in these mutants. It is unclear how an arginine kinase may influence the germ line, but its functions downstream of TOR signaling may be relevant. The antagonistic interactions between AMPK signaling and the TOR pathway have been described extensively [46–47], and it is widely accepted that AMPK negatively regulates TOR to block its growth enhancing effects during periods of energy stress [48]. Indeed, our own transcriptomic data confirms that the expression levels of many TOR components are increased in the AMPK mutant dauer larvae (S7A Fig). Curiously, ARGK-1 has also been shown to modulate AMPK, since active (S170 phosphorylated) AAK-2 was reduced in ARGK-1 and S6K double mutants [41]. From these findings, it seems probable that ARGK-1, AMPK and TOR do not function in a simple linear relationship, but rather there exists the possibility of feedback loops or some other form of regulation (S7B Fig).

**Fig 6.**
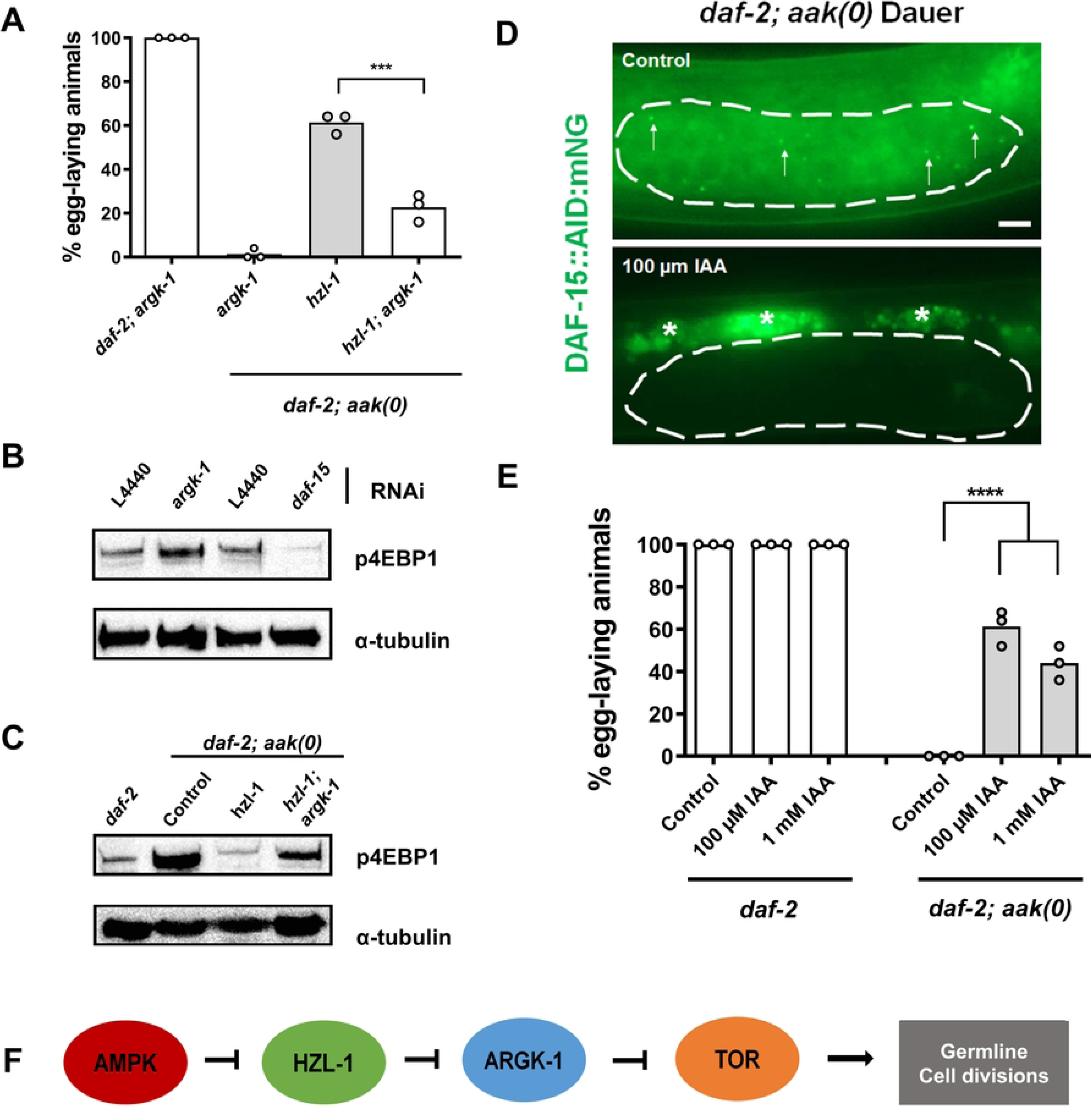
HZL-1 mRNA binding reverses *argk-1-*dependent modulation of TOR activity. (A) Post-dauer fertility of *hzl-1* and *argk-1* mutant strains. (B) and (C) Levels of P-h4EBP1 used as a proxy for TOR activity detected by Western blot in RNAi-treated animals or genetic mutants. Anti-P-h4EBP1 antibodies were used to detect phosphorylated h4EBP1 levels in animals. α-tubulin serves as a loading control (B) Dauer animals in *daf-;2 aak(0); hzl-1(0*) background treated with the indicated RNAi were used as samples. RNAi against *daf-15/*Raptor serves as a negative control. C) Dauer animals in indicated mutant backgrounds. Western analyses were performed on Day 2 dauer larvae for each experiment. Approximately ∼600 dauer larvae were used for each well, run on an 8% SDS-PAGE gel for 45 to 90 minutes, as needed. P-h4EBP1 and α-tubulin bands are from the same respective gels, with membranes cut and separated after membrane transfer for simultaneous antibody incubation. (D) Confocal micrograph images of *daf-2; aak(0)* animals with DAF-15::mNG::AID i.e. the DAF-15 degron. Animals grown on control plates (without auxin) (Top) express mNG in the germ line (within the white lines). DAF-15 localizes to discrete puncta, presumably lysosomes, as has been previously shown (white arrows). mNG signal is absent in animals exposed to 100 um IAA (Bottom). White asterisks denote non-specific signal from autofluorescence. Scale bar = 10 µm. (E) Post-dauer fertility of *daf-2* and *daf-2; aak(0)* animals with DAF-15:mNG::TIR and germline-expressed TIR following auxin treatment with either 100 µM or 1 mM IAA. Controls were grown on standard NGM plates without auxin during the dauer stage. Post-dauer fertility data represent the results from three independent trials, where the mean is represented by columns and values for individual trials indicated by small circles. n=50 for each trial. ****p < 0.0001 using one-way ANOVA for the indicated comparisons. (F) Model of AMPK-mediated regulation of HZL-1 and TOR. In the wild type, AMPK inhibits HZL-1, allowing for HZL-1-target mRNAs, such as *argk-1*, to promote germ cell quiescence. *argk-1* negatively regulates TOR, and when *argk-1* is bound by HZL-1, TOR activity is elevated. This drives the unscheduled germ cell divisions and subsequent hyperplasia observed in AMPK mutants during periods of energy challenge, resulting in post-dauer sterility.

**S7 Fig. Several TOR components have increased expression in AMPK dauer larvae compared to control**

(A) Heatmap of TOR component genes fold changes in indicated comparisons from RNA-Seq datasets. Heatmaps generated based on log2 fold change of indicated comparisons.

(B) Model of the TOR pathway highlighting a potential modulatory role of ARGK-1.

Our RNA-Seq data indicate that the expression of *argk-1* negatively correlates with the expression of the major TORC1 components in our mutants (Fig 5C, S7A Fig). It is therefore possible that ARGK-1 function indeed impinges on TOR activity. We monitored the effects of *argk-1* on a TOR activity sensor [49]. This *C. elegans* strain comprises a transgene expressing the human 4EBP1 protein, a well characterized target that is phosphorylated by TOR. An antibody raised against the TOR-phosphorylated form of the protein can then be used to detect the modification as a proxy for TOR activity. The sensor transgene was then crossed into our various mutant backgrounds, and TOR activity was assayed simply through Western blotting using the anti-P-h4EBP1 antibody.

We performed *argk-1* RNAi in the *aak(0); hzl-1* mutant strain and picked transgenic animals to use as samples for Western blot using the anti-P-h4EBP1 antibody. A stronger P-h4EBP1 signal was clearly visible in the sample from animals treated by the *argk-1* RNAi compared to those treated with the empty vector control (Fig 6B), suggesting that TOR activity is increased in response to the reduction of *argk-1* activity. To test the sensitivity of the sensor, we reduced TOR activity by eliminating the *C. elegans* orthologue of Raptor, *daf-15*, a key activating component in the TOR complex. As expected, we observed a weaker signal in animals treated with *daf-15* RNAi, indicating that the sensor faithfully reports TOR activity. Furthermore, the sensor also indicated that TOR activity was high in *aak(0)* dauer larvae, but lower in both *daf-2* control dauer larvae and *aak(0); hzl-1* mutants (Fig 6C). These findings correlate with our RNA-Seq data, which also mirrors this elevated expression of TOR components in *aak(0)* animals (S7A Fig). Together, these data support a model whereby the levels of *argk-1* are able to alter TOR activity, and HZL-1 binding to *argk-1* RNA and inhibiting it in turn leads to elevated levels of TOR.

### TOR is active in the germ line of AMPK mutant dauer larvae and contributes to post-dauer sterility

Our model suggests that when *argk-1* RNA is bound and inhibited by HZL-1, as is the case in *aak(0)* animals, this leads to elevated TOR activity. We and others have found that TOR contributes to aberrant germline proliferation during periods of starvation during the L1 diapause in AMPK mutants [46, 50], suggesting that a similar phenomenon may be occurring in the dauer stage as well.

To determine if misregulated TOR activation, specifically in the germ line of dauer *aak(0)* animals, influences the germline hyperplasia or has an impact on post-dauer fertility phenotypes in the AMPK mutants, we wanted to see how post-dauer fertility is affected when we inhibit TOR. We used an auxin-inducible degron (AID) system to circumvent the difficulties of using TOR mutants, which exhibit embryonic lethality or larval arrest phenotypes [18]. By fusing a degron sequence to DAF-15, a central component of TORC1 [51], we could effectively deplete TOR in various tissues as has been previously described [52–54]. Furthermore, we were able to take advantage of tissue-specific TIR expression to eliminate the protein in a spatially and temporally controlled manner.

The AID possessing transgenic animals were exposed to auxin during the dauer stage, and fertility was assayed in the post-dauer animals (protocol in S8A Fig). Expression of DAF-15, tagged to mNeonGreen, was largely absent in *daf-2* control dauer larvae, as expected. However, it was visible specifically in the germ line in *aak(0)* mutant animals (Fig 6D), consistent with our transcriptomic data. Expression in the germ line disappeared in animals exposed to auxin at either 100 µM or 1 mM, suggesting the degron system worked as intended. When assaying post-dauer fertility, we noted that loss of DAF-15 in the *daf-2* control background had no effect on fertility, but in the *aak(0)* animals, the addition of auxin improved fertility considerably, compared to controls (Fig 6E), suggesting elevated TOR activity in the germ line was contributing to post-dauer sterility of AMPK mutants.

**S8 Fig. HZL-1, ARGK-1 contribute to post-dauer sterility of AMPK mutants through germ line TOR regulation**

(A) Protocol for using an auxin-inducible DAF-15 degron to assess the role of TOR, downstream of AMPK/HZL-1/ARGK-1. *daf-2; aak(0)* animals with DAF-15::mNeonGreen::AID and TIR expressed under a soma- or germline-specific promoter were grown on NGM plates for 48 hours at 25°C, before being transferred to control or auxin supplemented plates, also at 25°C, degrade the protein specifically during the dauer stage. After 48 hours in dauer, some animals were picked for confocal imaging. The rest of the population was transferred to NGM plates and post-dauer fertility was assessed after they had recovered.

(B) Post-dauer fertility of *daf-2* and *daf-2; aak(0*) animals with DAF-15:mNG::TIR and soma-expressed TIR following auxin treatment with either 100 µM or 1 mM IAA.

(C) Confocal micrograph images of *aak(0); hzl-1* or *aak(0); hzl-1*; *argk-1* animals with DAF-15::mNG::AID. Animals grown on control plates (without auxin). Approximate region of germ line is shown between the white lines. DAF-15 enriched in puncta is also visible (white arrows). Asterisks denote non-specific signal from autofluorescence. Scale bar = 10 µm

(D) Post-dauer fertility of *aak(0); hzl-1* or *aak(0); hzl-1*; *argk-1* animals with DAF-15:mNG::TIR and germline**-**expressed TIR following auxin treatment with 100 µM IAA. Controls were grown on standard NGM plates without auxin during the dauer stage. Post-dauer fertility data represents three independent trials, with the mean represented by columns and values for individual trials indicated by small circles. n=50 for each trial. ***p < 0.001 using one-way ANOVA for the indicated comparisons.

There were no changes in post-dauer fertility when DAF-15 was degraded in the soma (S8B Fig), suggesting even if TOR is inappropriately active in somatic tissues, it has no negative impact on germline function in these animals. It should be noted, however, that it was difficult to perform this assay as soma-specific loss of DAF-15 induces developmental defects, including animals unable to escape dauer, or dying prematurely (also noted by [54]). Furthermore, we note that while TOR disruption can result in developmental defects in *C. elegans* (18), degradation of DAF-15 in AMPK mutants specifically in the dauer germ line did not result in developmental defects or arrest based on our observations.

We further tested our DAF-15 degron model in our *hzl-1* and *argk-1* mutants, and saw that the DAF-15 signal is absent in *aak(0); hzl-1* dauer larvae but visible in *aak(0); hzl-1; argk-1* mutants (S8C Fig), consistent with our measurements of TOR activity in these strains (Fig 6C). Similarly, there are no changes in post-dauer fertility when TOR is removed in the *aak(0); hzl-1* mutants, but the sterility seen in *aak(0); hzl-1; argk-1* mutants is partially suppressed by DAF-15 degradation (S8D Fig), suggesting the absence of ARGK-1 in this background once again leads to an upregulation of TOR activity that contributes to germ line defects. Our data thus indicate that unchecked TOR activity in the germ line in the dauer stage of *aak(0)* mutants is one contributor to the post-dauer sterility defects, and this is regulated by *argk-1*, a HZL-1 mRNA target.

Taken together, these data suggest that AMPK suppresses HZL-1 in the dauer stage most likely through its phosphorylation of S588, thereby ensuring quiescence. When AMPK is absent, HZL-1 is able to bind and inhibit RNAs such as *argk-1*, and low levels of *argk-1* result in elevated TOR activity, which contributes to the observed defects in the germ line, culminating in post-dauer sterility (Fig 6F).

## Discussion

The ability of organisms to respond to changes in environmental conditions is crucial for their survival and reproductive fitness. *C. elegans* can enter the dauer diapause when faced with environmental stressors, and to ensure the animals are fertile post-dauer, the germ cells must enter a quiescent state to preserve the integrity of the germ line throughout this stage. Previously, we have shown that the loss of the metabolic regulator AMPK results in germline hyperplasia and post-dauer sterility, among a host of somatic defects, as well as reduced dauer survival [17]. AMPK allows organisms to adjust to nutrient and energy stress, permitting cells/tissues to maintain homeostasis in the absence of sufficient nutrition. The kinase regulates a number of different targets, mostly through direct phosphorylation, and in its absence, the misregulation of these factors render the animals incapable of adjusting to the developmental and metabolic challenges of the dauer diapause.

Here, we have identified a mechanism that *C. elegans* employs to adjust to fluctuating metabolic requirements through its ability to regulate RNA metabolism. We investigated how a previously undescribed, putative RNA-binding helicase blocks the activity of key RNAs that otherwise promote a quiescent state in the *C. elegans* germ line, and how that protein is in turn regulated by the highly conserved metabolic regulator AMPK. By characterizing the various steps and identifying some of the molecular targets of this RNA helicase, we have simultaneously identified yet another means through which developmental plasticity can be fine-tuned, allowing for animals to adopt different metabolic and developmental states depending on the environments they are confronted with. It also further underscores the important role of RNA regulation as a major process by which organisms can control their metabolism, growth and development, and ultimately, their reproductive fitness.

All of our analyses indicate that this RNA-binding helicase, HZL-1, functions in the intestine in order to exert its control over the germ line. This type of cross-tissue communication has been previously described in *C. elegans* (55-56). Given the global transcriptomic changes and extensive remodeling that must occur during the dauer stage, it is perhaps not surprising that the animal responds to a localized input, in this case, nutrient status detected in the intestine, and is able to generate a response in other tissues, most importantly in the germ cells. In the case of HZL-1, the protein appears to target intestinal RNAs, such as *argk-1*, inhibiting them and their cognate genes in order to modulate downstream proliferation pathways, leading to growth (Fig 5). This likely occurs when animals are sufficiently fed and are on a trajectory towards reproductive development, whereas this pathway is suppressed in the dauer stage, when animals are typically quiescent. When activated aberrantly, as seen in AMPK-deficient dauer larvae, this results in extensive proliferation of the germ cells. This finding highlights the necessity for tight regulatory control by factors such as AMPK, as an otherwise beneficial pro-growth signal can rapidly become deleterious for the organism if misregulated. Our evidence suggests that the activation of AMPK leads to the phosphorylation of HZL-1 at a single residue in order to inhibit the protein when needed (Fig 2). This form of regulation by AMPK has been observed previously [27, 40, 57], although the precise mechanism by which phosphorylation results in the modulation of HZL-1 and its function remains to be elucidated. Based on protein structure predictions, we suspect that phosphorylation may induce a conformational change that suppresses the ability of the helicase to bind RNA (Fig 2D).

HZL-1 appears to function by binding mRNAs, thereby affecting their stability and likely preventing their translation (Fig 5C). While the precise mechanism by which it affects its targets remains unclear, there is evidence to suggest the protein behaves similar to other, well-characterized RNA helicases. One characteristic that lends credence to this theory is the existence of intrinsically disordered regions in the protein sequence of HZL-1. We demonstrated that removal of the first and third predicted IDR from HZL-1 not only disrupts the expression of the protein in the dauer stage, but also suppresses its function, as this modified version of the protein failed to rescue the mutant phenotypes in post-dauer animals (Fig 3). The formation of liquid-liquid condensates is often associated with these IDR domains [58], providing a biophysical niche for RNA-binding proteins to localize and interact with their cognate RNA targets [32]. Since several RNA helicases undergo phase separation in order to interface with RNAs or other RNA-binding proteins, we suspect HZL-1 may be doing the same. However, imaging of HZL-1, particularly in its endogenous form, proved to be difficult. The expression levels of the protein are likely too low to be observed by standard confocal microscopy. Conversely, intestinally-expressed HZL-1, while abundantly visible with an mKate tag, does not visibly form the discrete droplets that are typical of condensates [59]. However this does not exclude the possibility that these proteins can indeed associate into subsaturated macromolecular clusters that would not be visible by conventional light microscopy, as many other RNA-binding proteins do [60–61]. Further experiments will be required to confirm the biophysical properties of HZL-1, perhaps taking advantage of an *in vitro* model and/or quantitative light scattering microscopy.

The transcriptome of *aak(0); hzl-1* mutants differs dramatically from animals lacking AMPK alone. Indeed, the gene expression profile appears to be closer to wild-type, as many genes associated with reproduction and germline homeostasis are enriched in *aak(0)* dauer larvae, but are decreased in the *aak(0); hzl-1* mutant background (Fig 4C-D). This is consistent with previous findings that *aak(0)* dauer larvae skip the quiescence typical of dauer larvae and adopt a reproductive state instead. This leads to premature or discordant germ cell development. Indeed, the dauer transcriptome of *aak(0)* mutants is strikingly similar to the post-dauer transcriptome of wild-type animals (Fig 4A,D). In the absence of AMPK, *hzl-1* and other factors could contribute to this by activating growth pathways that ultimately result in widespread gene misregulation. As a result, when HZL-1 itself is disabled, there is a partial return to wild-type gene expression.

Our RNAi survey indicated that *parp-2* also regulates post-dauer sterility in AMPK mutants, similar to *hzl-1*. Epistasis analysis suggests that the two genes may function in the same pathway, or may act in parallel to converge on a common downstream effector(s), as there were no additive effects on fertility following RNAi of *parp-2* in a *hzl-1* deletion background (S1B Fig). *parp-2* was found to regulate PARylation in the germ line of PARG-1 mutants, which also exhibit reproductive defects [62]. We note high levels of both *parp-1* and *parp-2* mRNA in our *aak(0)* transcriptomic dataset, which are concomitantly reduced in *aak(0); hzl-1* animals and in *daf-2* control data sets. This pattern of expression is similar to the roughly 2000 reproductive/germ line genes that are misregulated in *aak(0)* and *hzl-1* mutants, all aberrantly expressed in *aak(0)* mutant dauer larvae, but the expression of which is corrected in *hzl-1* mutants. *parp-2* is thus likely to be one of numerous genes misregulated as a result of HZL-1 activity, resulting in premature activity in the germ line of dauer larvae and contributing to the disruption of germline integrity.

It was evident from the suppressor phenotype that HZL-1 contributed to both the developmental phenotypes and the observed changes in gene expression in the AMPK mutants. However, following the CLIP-Seq analysis, the primary mechanism through which HZL-1 regulate fertility in the AMPK mutants was ultimately revealed. Like many RNA helicases, HZL-1 binds RNAs and inhibits their function either through either through degradation or sequestration [44]. This hypothesis is supported by our observation that many mRNA targets of HZL-1 have reduced expression in *aak(0)* dauer larvae, but have increased expression in the *aak(0); hzl-1* mutants (Fig 5C).

It is still unclear if or how the HZL-1-bound targets exert an effect on dauer physiology. It is plausible that some are responsible for the distinct survival defect of the mutant [40] as several of them are implicated in metabolic processes (Fig 5A-B). We have previously identified a number of genes that, when compromised, suppress the reduced survival defect of *aak(0)* dauer larvae [63]. A large number of those genes are associated with metabolism. It is thus possible that some of the RNAs bound and inhibited by HZL-1 may in be regulators of those genes that contribute to the premature death of *aak(0)* dauer larvae, although no obvious candidate regulators stand out in our CLIP-Seq dataset. Notably, we detected RNAs bound to HZL-1 in both control and *aak(0)* backgrounds, although there were more RNAs bound in the AMPK mutants, where HZL-1 is presumably more active. It is unclear what function, if any, HZL-1 is carrying out in wild-type dauer animals.

We specifically identified *hmit-1.1* and *argk-1* as targets of HZL-1 that both impact on post-dauer sterility in the AMPK mutants (Fig 5C-D). It was surprising to see *hmit-1.1*, a metabolic transporter gene implicated in the *aak(0); hzl-1* post-dauer sterility phenotype. However, we suspect that this may be a result of the severe metabolic readjustments that occur in the absence of AMPK. While we have not previously tested the osmotic stress potential of AMPK dauer larvae, it is known that *C. elegans* dauer larvae can adapt to various abiotic stressors [64]. Furthermore, we have previously demonstrated that expression of *aak-2* in the excretory system in AMPK mutants suppresses the survival defect, and may also impact osmoregulation, which require the function of the excretory tissues [40, 65]. It is thus possible that AMPK mutant dauer larvae are unable to carry out an osmoprotective response as well as the wild-type in part due to a downregulation of *hmit-1.1*. Given that *C. elegans* have mechanisms to protect the germ line during periods of osmotic stress [66] it is similarly plausible that poor osmoregulation due to a lack of *hmit-1.1* leads to additional changes that affect the germ cells, thus contributing to the sterility of post-dauer AMPK animals.

*argk-1* stood out as the most enriched target bound to HZL-1 in *aak(0)* mutant dauer larvae. This arginine kinase is also the most enriched protein observed in S6K/*rsks-1* mutants in adult *C. elegans* based on mass spectrometry analysis [41], suggesting it is negatively regulated by *rsks-1* and thus acts downstream of, or in parallel, to TOR signaling [67–68]. Curiously, *argk-1* was also shown to be necessary for the phosphorylation and activation of AAK-2 in the *rsks-1* mutant background, a process that is required for the increased longevity of the *rsks-1* animals. We revealed that the loss of *argk-1* reverts the suppression of post-dauer sterility in *hzl-1* mutants (Fig 6A). It came as a surprise that the protein could have such a distinct role in reproductive development, given its function in other species, as creatine kinase has been more directly linked to energy modulation [69]. Previous analyses demonstrated that ARGK-1 was associated with TOR complex targets, although the relationship between ARGK-1 and TOR were not clear [41]. Our RNA-Seq data suggested high TOR activity was a phenotype of AMPK dauer larvae (S7A Fig), and we confirmed a role of TOR regulation through the use of a TOR activity sensor, with which we saw an increase in TOR activity following a knockdown of *argk-1* (Fig 6B), suggesting ARGK-1 is a negative regulator of TOR in this developmental context.

Nevertheless, it remains unclear how ARGK-1 is able to influence TOR activity, specifically in the germ line, but there is evidence that this regulation may be indirect. In mammals, creatine kinase, the closest ortholog of ARGK-1, is an enzyme that serves as an ATP buffer [70]. The ability of creatine kinase to affect metabolism may be conserved in *C. elegans*. If that is the case, it is perhaps not surprising that TOR activity, which fluctuates in response to various metabolic changes, is also affected by the activity of ARGK-1. Indeed, studies have demonstrated that changing the levels of creatine kinase can have an impact on TOR activity. Specifically, overexpression of the creatine kinase brain (Ckb) isoform in cells downregulates mTOR [71], consistent with our findings of ARGK-1 being a negative TOR regulator. Conversely, another study demonstrated that loss of Ckb reduced CD8^+^ T cell expansion because of weakened mTOR signaling [72]. The complex relationship between creatine kinase activity and the TOR pathway is likely indicative of different metabolic responses that are required in specific tissues or other conditions. There are notable physiological differences between these mammalian cell models and the *C. elegans* dauer model we have examined, and it is also not well established at the biochemical level how similar the ARGK-1 and mammalian creatine kinase might actually be. Nevertheless, there is established precedent that these enzymes can affect the action of the TOR pathway, and further metabolomic analyses could potentially clarify the specific function of ARGK-1 and its role in metabolic regulation.

The role of the TOR complex in regulating longevity, stress resistance, and other related phenotypes have been thoroughly studied, particularly in *C. elegans* [18, 67,73–75]. However, it also plays a role in maintaining fertility, in nematode species and other models [53,76]. Our work provides yet another example of how TOR activity must be tightly controlled, specifically in a period of acute metabolic stress, in order to preserve reproductive fitness. Previously, it has been demonstrated that increased TOR activity contributes to aberrant germline proliferation in L1-starved *aak(0)* animals [46]. Our degron experiment, where we depleted TOR specifically in the germ line of dauer *aak(0)* mutants, also indicated that TOR may function in a similar manner in the dauer stage (Fig 6).

Regulation of TOR by AMPK has been well-studied [26, 77], and so it is not surprising that TOR activity contributes to post-dauer phenotypes in AMPK-deficient mutants. TOR is one of several metabolic regulators which controls the growth of organisms, and thus must be modulated during the quiescent dauer phase. Without AMPK serving as a master regulator, however, TOR activity is elevated, leading to deleterious developmental defects as the animal can no longer reconcile its continued growth with the nutritional limitations of the dauer stage. This revelation highlights the importance of coordinating development with metabolic status. Animals enter the dauer stage when faced with harsh environmental conditions, such as a lack of nutrients, and a subsequent downregulation of developmental programs is critical such that the meagre resources available are not used inappropriately, and the organism can instead survive until conditions improve.

Our work identifies a throughline between the metabolic regulator AMPK and the maintenance of germline quiescence in the dauer stage through the inhibition of TOR signaling (Fig 6F). Rather than direct regulation, however, we identify a potentially rapidly reversible mechanism by which HZL-1, a putative RNA helicase, modulates levels of various RNAs in the intestine in the absence of AMPK, resulting in widespread changes to the transcriptome, and activation of downstream signaling pathways, including TOR. Specifically, the inhibition of a metabolic gene, *argk-1*, contributes to the disruption of germline quiescence. Our unbiased genetic analysis of *hzl-1* has consequently expanded our understanding of the TOR signaling pathway. We demonstrated how the absence of regulators such as AMPK lead to strong TOR expression in the germ line even in the quiescent dauer stage, and how this directly impacts development and fertility. Most notably, we highlighted the role of a novel negative regulator of TOR in *C. elegans*, ARGK-1, although the precise mechanism through which this regulation might be executed remains to be fully elucidated.

## Materials and methods

### Maintenance of *C. elegans* strains

Animals were grown on NGM plates seeded with *E. coli* OP50. *C. elegans* strains with the *daf-2(e1370)* mutation were maintained at 15 °C to ensure no dauer formation. Synchronization and sterilization of *C. elegans* embryos was performed using sodium hydroxide and sodium hypochlorite solutions, using standard protocols [78]. For most experimental procedures, animals were bleach-synchronized and allowed to hatch into L1 larvae before being dispensed onto plates, in order to ensure all animals were of the same developmental stage. Transgenic animals were maintained by picking rollers

List of strains used is listed in S1 Table.

### Proteomic analysis

Putative AMPK phosphorylation sites were identified using GPS 6.0 [79]. *C. elegans* proteome sequences were inputted into the program, and putative p-sites were identified using the species-specific batch predictor module, using the high threshold setting. Putative targets were further screened based on relevance to RNA pathways or functions.

Alphafold 3 was used to predict and analyze protein structures [29].

### RNA interference experiments

RNAi experiments were performed using feeding, as described [80]. Briefly, dsRNA producing bacteria was grown with Ampicillin selection overnight at 37 °C using LB, then seeded onto NGM plates containing 1 mM IPTG and 50 µg/mL Ampicillin (i.e. ‘RNAi plates’) the following day. Plates were grown for a minimum of one day at room temperature before the addition of animals. For all RNAi experiments, L1 synchronized animals were put onto plates, then moved immediately to 25 °C to promote entry into dauer.

### Post-dauer fertility assay

Animals were kept at 25 °C for 4 days following L1 synchronization, then moved to 15 °C for further observation. Animals were picked onto either NGM plates, or plates growing RNAi identical to the plate being picked from, depending on the experiment in question. Singled animals were allowed develop for a minimum of one week, after which fertility was assayed by checking for the presence of F1 progeny. Three replicates were done for every post-dauer fertility assay.

### Confocal microscopy

2% agarose microscopy pads were created on microscope slides, and 20 mM levamisole (500 mM when mounting dauer animals) was deposited onto cooled agarose. Animals were picked onto slides and allowed to sit for approximately five minutes to induce paralysis, before being covered by a slide cover. Edges of slide covers was sealed using nail polish as needed. Slides were viewed on a Leica DMI 6000B inverted microscope equipped with a Quorum WaveFX spinning Disc and EM CCD, under various magnifications with and without oil immersion. Low-resolution images were obtained through a single stack, while high resolution images were taken through a Z stack with a range of 10-5 µm and intervals of 0.2 µm. Images were processed via deconvolution and maximum projection for the Z stack, and further processed as needed using the application Fiji.

### Western Blot

Western Blotting was performed using standard protocol with 6-15% SDS-Page gels, as needed. 20ul of collected worm samples in PBS, typically ∼600 dauer larvae per sample, were loaded into wells and run at 85V for the stacking gel, then at 100-150V for the resolving gel, as required. Following membrane transfer, membranes were cut based on protein size i.e. a single membrane might be cut in half at the 100 kDa mark in order to obtain one membrane containing bands at 50 kDa and another with bands at150 kDa. Membranes were then incubated with either anti-GFP, anti-α-tubulin, or anti-P-h4eBP1 antibodies [49]. After overnight primary antibody incubation, membranes were washed with PBST 3 times, 15 minutes each, followed by secondary antibody incubation for 2 hours. After another 3 PBST washes, membranes were soaked with total 1 ml Clarity Western ECL Substrate, then imaged using the MicroChemi (DNR Bio Imaging Systems) and GelCapture software (Version 7.0.18), using exposure times of 0.5s to 30s, as required.

### Generation of transgenic lines

High fidelity PCR was performed to amplify gene fragments of interest from *C. elegans* worm lysis, as well as to linearize relevant plasmid vectors. Constructs were generated through Gibson assembly [81], and following transformation into competent cells, cracking was carried out and gel electrophoresis performed to identify colonies with plasmid DNA of the correct size.

Plasmids were microinjected into *C. elegans* strains of interest along with a *rol-6* marker for selection. Concentrations of ∼30 µl was used for the selection marker and ∼120-160 µl for the plasmid of interest, diluted in ultrapure water. F2 transformants were selected by picking rollers and maintained as independent lines.

### RNA extraction and sequencing

For RNA extractions, animals were washed with M9 and pelleted into Eppendorf tubes. Standard Trizol extraction [82] was performed on relevant samples to extract any present RNA.

Briefly, 200 µl of Trizol was added to 20 µl of packed worms, and tubes were subjected to freeze thawing with liquid nitrogen at room temperature. After four freeze-thaws, samples were kept at 4 °C for approximately 1 hour to settle particulates, then briefly centrifuged at high speeds. The solution from each tube was then transferred to a second set of tubes, without disturbing the pellet. 50 µl of chloroform was added to the second set of tubes, followed by 30 seconds of agitation by vortex mixer, 3 minutes of rest, then 15 minutes of centrifugation at high speeds. The aqueous phase from each tube was then isolated and placed into another set of tubes, at which point the chloroform extraction was repeated.

Following the second extraction of the aqueous layer, 125 µl of isopropanol was added to samples and then tubes were inverted, followed by several minutes rest, and then 10 minutes of centrifugation at high speeds. The liquid from tubes was aspirated without disturbing the pellet, and then samples were washed with 500-1000 µl of 70% ethanol diluted in ultrapure water. After another 5 minute centrifugation, solutions were aspirated as much as possible, and then tubes were air dried over the course of one hour, or as needed. Samples were dissolved in 10-20 µl ultrapure water at room temperature, and their approximate concentration measured using a Nanodrop. All centrifugation was carried out at 4 °C.

Following purification, total RNA from each sample was run on 1% agarose gel to confirm presence of 18s and 28s ribosomal RNA bands, confirming integrity. RNA was stored for <3 days at -80 °C before sending off for sequencing. Sequencing was performed using NextSeq Midoutput with 2x75bp coverage.

### Analysis of RNA-Seq data

RNA-Seq data was analyzed using DESeq2. Volcano plots of RNAseq data were created using a custom script in R Studio. Gene ontology was carried out using WormBase’s Gene Set Enrichment Analysis. Bubble plots, heatmaps, and Venn diagrams were generated using custom R scripts, Microsoft Powerpoint, or online resources (https://www.bioinformatics.com.cn, [83]). Data analysis, thresholding, pairwise comparisons etc. were performed using Microsoft Excel.

### Cross-linking immunoprecipitation

Large quantities of dauer worms from relevant strains were collected through washing with M9, and then were cross-linked with formaldehyde before being frozen at -80 °C. Samples were thawed and lysed using a sonicator, followed by immunoprecipitation.

### Preparation of beads

Immunoprecipitation was carried out using 25 mg Protein A beads per sample. 1 ml PBS was added to each sample of beads and allowed to swell for 1 hour, followed by brief centrifugation at low speeds, after which liquid was aspirated. Dilution buffer (PBS + 1% BSA) was added to beads in a 1:1 ratio and tubes were rotated at 4 °C for 10 minutes. After centrifugation and aspiration of liquid, 12 µl of relevant antibody serum (GFP or FLAG) was added to each sample, followed by 1 hour rotation at 4 °C. Afterwards, beads were aspirated followed by another dilution buffer incubation and wash as before, followed by a PBS wash.

Then, freshly made DMP, created from 13 mg/ml DMP stock in 1 ml of wash buffer (0.2M triethanolamine in PBS) was added to tubes at a 1:1 ratio, followed by 30 minutes room temperature incubation. Beads were then washed with wash buffer with 5 minutes of rotation at room temperature before aspiration. DMP treatment and incubation was then repeated twice, with washing as before. After the 3^rd^ treatment, quenching buffer (50 mM ethanolamine in PBS) was added to samples in a 1:1 ratio followed by 5 minutes of rotation at room temperature and then centrifugation and aspiration. Samples were then washed with elution buffer (1M glycine) with 10 minutes rotation at room temperature, and this step was then repeated. Finally, samples were washed with PBS + 0.1% Tween 3 times. At this point, beads were incubated with lysed samples as needed, rotating at 4 °C overnight.

The following day, all samples were centrifuged and aspirated, followed by 3 PBS washes. At this point, beads were aliquoted and used directly for Trizol RNA extraction (as described above).

### AID experiments

Auxin-supplemented NGM plates were created through addition of indole-3-acetic acid (IAA) from a 1 M stock solution to molten agar solutions once agar was sufficiently cooled. IAA was added to create plates with a working concentration of either 100 µM or 1 mM.

For AID experiments, animals were synchronized and grown on standard NGM plates, then picked onto Auxin plates for 48 hours at 25 °C, as described (S8A Fig), before being transferred to NGM plates for growth at 15 °C. As a negative control, degron-tagged animals were also grown on standard NGM plates without auxin. Depletion of the degron-tagged protein of interest was confirmed using confocal microscopy. Confocal imaging of animals from auxin plates, as well as post-dauer fertility assays, were performed as described above.

### RT-qPCR

Trizol RNA extraction was performed on washed worm samples, as described above. Approximately 1 µg of extracted total RNA was converted to cDNA using the High-Capacity RNA-to-cDNATM Kit (AppliedBiosystems, 4387406). The 2x SyberGreen qPCR Master-mix (ZmTech Scientifique, Q2100N) was used for qPCR using 1-2 ng of cDNA. A Bio-Rad CFX384 Real-Time 96-well PCR qPCR Detection System (Bio-Rad) coupled with the CFX Maestro Software (Bio-Rad) was used to analyze qPCR data. *tba-2* served as a refence gene for calculations. 3 replicates were performed for each qPCR reaction, with 1 additional replicate using DNAase-free H_2_O instead of cDNA. Fold change calculations were performed using the delta Ct method.

For CLIP-qPCR, immunoprecipitation was performed on relevant samples using both anti-GFP and anti-FLAG (i.e. a non-specific antibody). For qPCR, both antibody samples as well as the original input lysate was subjected to Trizol RNA extraction. For calculations, the “% Input% method was used. Briefly, the dilution factor of the original lysate (before IP) was taken into account (specifically, a dilution factor of 50), and relative expression was calculated compared to the lysate. The delta Ct method was used, whereby the adjusted Ct of the input was calculated by subtracting the Log_2_ of the dilatation factor from the Ct of each input. The delta Ct was calculated by subtracting the GFP and FLAG Ct values for each sample from the corresponding adjusted input Ct. Finally, % input was calculated as 2^-(adjusted Ct) for both the GFP and FLAG Ct values for each sample.

